# Molecular population genetics of *Sex-lethal* (*Sxl*) in the *D. melanogaster* species group - a locus that genetically interacts with *Wolbachia pipientis* in *Drosophila melanogaster*

**DOI:** 10.1101/2021.01.10.426102

**Authors:** Vanessa L. Bauer DuMont, Simone L. White, Daniel Zinshteyn, Charles F. Aquadro

## Abstract

*Sex-lethal* (*Sxl)* is the sex determination switch in *Drosophila*, and also plays a critical role in germ-line stem cell (GSC) daughter differentiation in *Drosophila melanogaster*. Three female-sterile alleles at *Sxl* in *Drosophila melanogaster* were previously shown to genetically interact to varying degrees with the maternally inherited endosymbiont *Wolbachia pipientis.* Given this genetic interaction and *W. pipientis’* ability to manipulate reproduction in *Drosophila*, we carried out a careful study of both the population genetics (within four *Drosophila* species) and molecular evolutionary analysis (across 20 *Drosophila* species) of *Sxl*. Consistent with earlier studies, we find that selective constraint has played a prominent role in *Sxl’s* molecular evolution within *Drosophila*, but we also observe patterns that suggest both episodic bursts of protein evolution and recent positive selection at *Sxl*. The episodic nature of *Sxl’s* protein evolution is discussed in light of its genetic interaction with *W. pipientis*.

## Introduction

Reproductive success is a key fitness trait governed by a plethora of gene regulatory networks. Despite the presumed functional constraint for such critical genes, many of them have been shown to be evolving rapidly due to positive selection at the amino acid level (Clark et al. 2006; Chapman 2008; Wong and Rundle 2013; Popovic et al. 2014). For some sets of reproductive loci this observation makes intuitive sense, including loci involved in species-specific gamete recognition and those involved in coevolutionary conflict between the sexes. Interestingly, non-neutral patterns of amino acid evolution have also been detected at loci involved in the differentiation of germ-line stem cells (GSCs) (Civetta et al. 2006; Bauer DuMont et al. 2007; Langley et al. 2012; Pool et al. 2012; Choi and Aquadro 2015; Flores et al. 2015a). The temporal and spatial expression of these GSC regulating loci does not coincide with that expected for genes influenced by sperm competition, sexual selection, inbreeding avoidance and gamete recognition (reviewed in Clark et al. 2006).

Several GSC regulating loci have functions outside the germline (e.g., Bell et al. 1988; Yi et al. 2008; Saito et al. 2010; Le Thomas et al. 2013) so their signatures of positive selection could be due to non-gametogenic functions. For many others it is possible that the positive selection is acting directly on gametogenic functions. We have previously hypothesized that interactions with maternally transmitted endosymbionts could be a gametogenesis-specific driver of positive selection (Bauer DuMont et al. 2007; Flores et al. 2015a and 2015b). One such endosymbiont, *Wolbachia pipientis,* infects an estimated 66% of arthropod species (Hilgenboecker et al. 2008) and is an obligate maternally transmitted endosymbiont that has been shown to manipulate host reproduction systems (Werren et al. 2008). *Wolbachia pipientis* infection has also been shown to have beneficial consequences. For example, infection in *Drosophila melanogaster* conveys greater resistance to viruses (Hedges et al. 2008; Teixeira et al. 2008; Chrostek et al. 2013) and has been shown to increase female fecundity on low and high iron diets relative to uninfected flies (Brownlie et al. 2009).

Here we explore the phylogenetic patterns of positive selection for amino acid diversification at *Sxl*, the sex determination master switch in *Drosophila* development (Bell et al. 1988) that also plays a critical role, along with *bag of marbles* (*bam*), in the maturation of cystoblasts during oogenesis (Chau et al. 2009). Proper *bam* function is essential for the start of cystoblast differentiation (McKearin and Spradling 1990). Epistasis experiments revealed that *bam* requires *Sxl* activity for proper germline stem cell daughter differentiation and the presence of both proteins is proposed to be responsible for regulating sex-specific gametogenesis (Chau et al. 2009; Chau et al. 2012; Shapiro-Kulnane et al. 2015).

In addition to genetically interacting with one another, hypomorphic alleles of both proteins have been found to genetically interact with *W. pipientis* infection. Starr and Cline (2002) reported that *W. pipientis* infection partially rescues the female-sterile phenotype caused by three *Sxl* alleles, though to differing degrees for each allele. Similarly, *W. pipientis* infection can mitigate the mutant phenotype of a hypomorphic allelic combination at the *bam* locus (Flores et al. 2015b). Starr and Cline (2002) also reported that *W. pipientis* infection did not rescue mutants in three other genes with ovarian tumor phenotypes (*snf*^*1621*^, *otu*^*11*^, and *mei-P26*^*fs1*^). These results suggest a specific, possibly physical, interaction of *W. pipientis* or its gene products with the *Sxl* and *bam* genes or gene products.

Using methods that consider both polymorphism and divergence we and others have previously shown that in *D. melanogaster* and *D. simulans,* the *bam* locus is evolving rapidly at the amino acid level due to positive selection (Civetta et al. 2006; Bauer DuMont et al. 2007). Phylogenetic methods for detecting selection across a number of *Drosophila* species did not detect evidence of positive selection, suggesting the selection pressures acting on *bam* are episodic. On the other hand, recent studies of the molecular evolution of *Sxl* have focused on the evolution of *Sxl’s* role as the sex determination master switch in *Drosophila* development (Mullon et al. 2012; Zhang et al. 2013). Mullon et al. (2012) used the maximum likelihood based phylogenetic methods implemented in PAML (Yang 2007) within and between 3 families of Diptera (Drosophilidae, Tephritidae and Muscidae), and detected both a relaxation of functional constraint and positive selection acting across amino acid sites along the lineage leading to *Drosophila*. However, they observe no evidence of positive selection among the 12 *Drosophila* species suggesting that the positive selection they observed was associated with the acquisition of the sex determination function in the common ancestor of *Drosophila* species.

Here we test for evidence departures from selective neutrality at *Sxl* within the genus *Drosophila* specifically by incorporating sequence polymorphism and divergence data, and test for departures consistent with lineage specific positive selection. First, we obtained high-quality Sanger sequencing polymorphism data at *Sxl* in the following species: *Drosophila melanogaster, D. simulans, D. ananassae* and *D. pseudoobscura*. Except for *D. pseudoobscura,* these species are currently infected with *W. pipientis* (Mateos et al. 2006). Second, we searched 8 additional sequenced *Drosophila* genomes to annotate *Sxl* orthologs, bringing the total number of *Drosophila Sxl* orthologous sequences to 20 for phylogenetic analysis. The inclusion of polymorphism data and additional *Drosophila* species allowed us greater power to detect potential signatures of positive selection at *Sxl* within the genus *Drosophila*.

## Materials and Methods

All DNA was extracted using the Qiagen puregene kit A DNA isolation kits (Qiagen). For *D. melanogaster,* we sequenced *Sxl* in 20 extracted X-chromosome lines from Zambia Africa, which was made through a series of crosses using the balancer X chromosome line Fm7a and isofemale Zambia lines (Pool et al. 2012). For *D. simulans*, we used 10 lines from a Madagascar population sample collected in 1998 by J.W.O. Ballard. For *D. pseudoobscura* we included 29 lines collected from Mesa Verde National Park, Kaibab National Park and the Bosque Del Apache National Wildlife Refuge provided by Stephen Schaeffer. Our line of *D. miranda* was provided by Doris Bachtrog. Finally, we also surveyed *Sxl* variation across 12 lines of *D. ananassae* collected from Bangkok, Thailand provided by Wolfgang Stephan, and a single line of *D. bipectinata* from the UCSD Species Stock Center (stock 0000-1029.01). We also sequenced a single *Sxl* allele from *D. guanche* (obtained from the *Drosophila* Species Stock Center at University of California San Diego; stock number 14011-0095.01), and *D. atripex* (provided by Artyom Kopp), which were used for analyzes that require divergence for the *D. pseudoobscura* and *D. ananassae* datasets respectively. Our sequences are available via Genbank accession numbers KT935592 - KT935663. All other sequences used in our analyses were those from *Drosophila* 12 Genomes Consortium et al. (2007) or Chen et al. (2014) and downloaded from Flybase.

While there are multiple *Sxl* transcripts, due to alternative splicing, most of them include exons 5 through 8 in *D. melanogaster* according to Flybase genome annotations (St Pierre et al. 2014). We PCR amplified and sequenced exons 5 through 8. We used Promega GoTaq for amplification following their standard protocol. Sanger sequencing was performed using the PCR primers and internal sequencing primers (primer sequences available upon request) through the Cornell Institute of Biotechnology Genomics Facility (https://www.biotech.cornell.edu/core-facilities-brc/facilities/genomics-facility).

The following triplets of species were independently aligned using MegAlign (DNASTAR Inc., Madison WI): *D. melanogaster, D. simulans, D. yakuba; D. ananassae, D. atripex*, *D. bipectinata*; and *D. pseudoobscura*, *D. miranda, D. guanche.* Gaps within coding regions were manually adjusted to ensure the sequences remained in-frame. Population genetic analyses were performed using DNAsp 5.10.1 (Librado and Rozas 2009). When using coalescent simulations to obtain the *P*-values of site frequency based tests we incorporated recombination rate estimates (following Przeworski et al. 2001). For the *D. melanogaster* and *D. simulans* datasets, we used the *Drosophila melanogaster* Recombination Rate Calculator (Comeron et al. 2012) estimate of r = 3.34×10^−8^ recombinants per base-pair per generation for the *Sxl* region of the X chromosome. This translates to an estimated R = 152 for the 1500 base pair region sequenced (R = 3N_e_r since *Sxl* is X-linked and assuming a population size of 1.0×10^6^). For *D. ananassae* and *D. pseudoobscura* the values of R were estimated from the polymorphism data directly using DNAsp 5.10.1 (Librado and Rozas 2009) as R=194 and R=6, respectively.

For *D. melanogaster* we also incorporated demography into our neutral simulations when obtaining our significance cut-offs. These simulations were done using the program msABC (Pavlidis et al. 2010). There is growing evidence that African populations of *D. melanogaster* have experienced changes in effective population size over time (Glinka et al 2003; Li and Stephan 2006; Hutter et al 2007; Haddrill et al 2008; Duchen et al 2013; Singh et al 2013). However, given the large effective population size of these species and signatures of a high rate of adaptation (e.g., Begun et al 2007; Langley et al 2012), inferring demographic parameters is challenging. Because of this we simulated three different scenarios: standard neutral equilibrium model, standard neutral with exponential growth as estimated by Hutter et al (2007), and standard neutral with a 3 phase bottleneck as estimate by Duchen et al. (2013). We supplied msABC with uniform prior distributions for theta and all demographic parameters. The prior distribution for theta for *D. melanogaster* was obtained from Pool et al. (2012) and ranged between 0.006 and 0.009 per site. The resulting *P*-values are the proportion of simulated datasets that were less than (for negative statistics) or greater than (for positive statistics) our observed test statistic for *Sxl*. The *P*-values were adjusted for multiple testing following the Bonferroni method.

The McDonald-Kreitman test (MKT) was done manually following the method’s original implementation (McDonald and Kreitman 1991) by combining polymorphism from multiple species if it was available. If a position in the alignment had more than one nucleotide segregating within a species’ population sample, it was labeled as polymorphic. Divergent sites were those for which all alleles from one species differed from all the alleles of the other two species.

To test for evidence of departures from neutrality for synonymous sites at *Sxl*, we used the method of Bauer DuMont et al. (2005). This method looks for differences in the rates of preferred and unpreferred codon substitutions per site in a manner similar to a dN/dS comparison (Nei and Gojobori 1986). Statistical significance is assessed by a 2×2 contingency table comparison.

In order to test for evidence of departures from selective neutrality in rates and patterns of sequence evolution at *Sxl* across a broader group of *Drosophila* species, we first retrieved the *Sxl* gene region sequences from FlyBase for the following 20 *Drosophila* species: *D. melanogaster, D. sechellia, D. yakuba, D. erecta, D. eugracilis, D. ficusphila, D. rhopaloa, D. elegans, D. takahashii, D. biarmipes, D. kikkawae, D. bipectinata, D. ananassae, D. miranda, D. pseudoobsura, D. persimilis, D. willistoni, D. mojavensis, D. virilis, and D. grimshawi*. *Sxl* is an alternatively spliced locus. To ensure we are analyzing orthologous exons, we first made alignments of the entire gene region (introns and exons) using the web-based versions of the alignment programs Muscle (Edgar 2004) (http://www.ebi.ac.uk/Tools/msa/muscle/). We then used the annotated exons of *D. melanogaster* as a guide to identify orthologous coding sequences (CDS) for *Sxl’s* Isoform L (6 exons - the female specific splice variant; Bell et al. 1988) from each aligned sequence. This isoform contains the poly-proline region of the Sxl protein where the female-sterile *Sxl* variants are located (Starr and Cline 2002).

We estimated a *Sxl* gene tree across these 20 *Drosophila* species using Mega 5.1 (Tamura et al. 2011). A maximum likelihood tree was estimated using all nucleotide sites, default parameters, and the GTR substitution model with gamma distributed site variation. To test for selection across the estimated *Sxl* tree we used Hyphy (Kosakovsky Pond et al. 2005) run online using the DataMonkey website (http://www.datamonkey.org/). A model selection procedure was conducted to determine that the best nucleotide substitution model for the data was TrN93 (Tamura and Nei 1993) which was used in all subsequent analyzes. We ran GARD (Kosakovsky Pond et al. 2006a; Kosakovsky Pond et al. 2006b) to look for evidence of recombination across these species at *Sxl* (that would reflect incomplete ancestral polymorphism sorting) using a general discrete site-to-site rate variation and 3 rate classes. To detect evidence of purifying and/or diversifying selection across sites and lineages we used the following Hyphy programs: BranchRel, GAbranch, FUBAR and MEME.

## Results

We surveyed DNA variability at the population level at *Sxl* for four species of *Drosophila* (*D. melanogaster, D. simulans, D. ananassae* and *D. pseudoobscura*), the first three of which have evidence of current *W. pipientis* infection (Mateos et al. 2006). Even though 37 lines of *D. pseudoobscura* were surveyed for infection, this species appears to be currently uninfected with *W. pipientis* (Mateos et al. 2006).

We find that Sxl is a very conserved protein with little to no nonsynonymous polymorphism or divergence within or between the 20 *Drosophila* species surveyed (Table 1). In addition, levels of synonymous polymorphism and divergence between *D. melanogaster* and *D. simulans* are below the average reported by Andolfatto (2005). The same is true for synonymous variation observed within and between *D. pseudoobscura* and *D. miranda* as compared to that reported by Haddrill et al. (2010). While synonymous polymorphism is slightly lower at *Sxl* within *D. ananassae* the level of divergence between *D. ananassae* and *D. atripex* is similar to values previously reported (Grath et al. 2009; Choi and Aquadro 2014).

**Table 1.** Levels of within species variation and divergence between *Drosophila* species and results of tests to detect departures from a neutral site frequency spectrum

| | $\theta$ | $\pi$ | Div<sup>c</sup> | Taj-D<sup>a</sup><br>(*P*-value<sup>b</sup>) | FW-H<sup>a</sup><br>(*P*-value<sup>b</sup>) |
| --- | --- | --- | --- | --- | --- |
| <u>*D. melanogaster* (n = 20)</u> |  |  |  |  |  |
| Synonymous | 0.012 | 0.011 | 0.069 | -0.267 | -2.26 |
| Nonsynonymous | 0 | 0 | 0 | (0.214; 0.80; 0.70) | (**<0.0002**; **<0.0002**; **<0.0002**) |
| Intron | 0.019 | 0.019 | 0.083 |  |  |
| <u>*D. simulans* (n = 10)</u> |  |  |  |  |  |
| Synonymous | 0.017 | 0.013 | 0.069 | -1.045 | 3.47 |
| Nonsynonymous | 0 | 0 | 0 | (**0.001**) | (0.87) |
| Intron | 0.031 | 0.025 | 0.083 |  |  |
| <u>*D. ananassae* (n = 13)</u> |  |  |  |  |  |
| Synonymous | 0.010 | 0.010 | 0.209 | 0.349 | -4.333 |
| Nonsynonymous | 0.000 | 0.000 | 0.003 | (0.166) | (0.038) |
| Intron | 0.021 | 0.025 | 0.299 |  |  |
| <u>*D. pseudoobscura* (n = 29)</u> |  |  |  |  |  |
| Synonymous | 0.011 | 0.007 | 0.073 | -1.293 | -0.539 |
| Nonsynonymous | 0.0004 | 0.0001 | 0.002 | (0.038) | (0.311) |
| Intron | 0.014 | 0.008 | 0.111 |  |  |

<sup>a</sup> test statistic when using all sites in the analysis: Tajima's D (Taj-D) and Fay and Wu's H (FW-H)
<sup>b</sup> Proportion of simulated datasets that were equal to or less than (for negative statistics) or equal to or greater than (for positive statistics) our observed test statistic for *Sxl*. These are two-sided tests, for which we do two tests per species resulting in a significant cut-off level of 0.0125 (0.025/2). For *D. melanogaster* the *P*-value was calculated by simulating different demographic scenarios listed in this order in parenthesis: Standard neutral; Bottleneck with exponential growth and a 3 Epoch bottleneck (as described in Materials and Methods). *P*-values significant after multiple testing corrections in bold
<sup>c</sup> Uncorrected divergence between the following species pairs: *D. melanogaster* and *D. simulans*; *D. ananassae* and *D. atripex*; and *D. pseudoobscura* and *D. miranda*

Previous studies have reported a skew in the Site Frequency Spectrum (SFS) toward rare alleles, as illustrated by a general negative Tajima *D* (Tajima 1989) test statistic, in all of the species included in our study (Kliman et al. 2000; Machado et al. 2002; Das et al. 2004; Andolfatto 2007; Grath et al. 2009; Haddrill et al. 2010; Jensen and Bachtrog 2011;). At *Sxl,* we observe a negative Tajima *D* for all species, except *D. ananassae*. Tajima *D* in *D. simulans* rejects the hypothesis of a SFS at equilibrium with 67% (33/49) of the polymorphisms being singletons. These singletons are evenly distributed across synonymous (sevem singleton/10 total) and intron (26 singleton/39 total) sites. At *Sxl,* Fay and Wu’s *H* is negative in all species but *D. simulans*. Fay and Wu’s *H* statistic is significantly negative in *D. melanogaster* even when considering two different demographic scenarios estimated for African populations of this species (Hutter et al. 2007; Duchen et al. 2013).

The McDonald-Kreitman Test (MKT; McDonald and Kreitman 1991) is used to detect departures from the neutral expectation that synonymous and nonsynonymous variants will have similar ratios of within to between species variation. A rejection in the direction of an excess of nonsynonymous divergence is typically interpreted as evidence of repeated amino acid substitutions due to positive selection. As seen in Table 1 there are few nonsynonymous changes at *Sxl*. The MKT does not reject neutral expectations when *D. ananassae* and *D pseudoobscura* polymorphism is compared to divergence to *D. atripex* and *D. miranda*, respectively (Table 2). We observe no amino acid differences within or between *D. melanogaster* and *D. simulans*. However, when considering the *Sxl* sequence from two additional and closely related Drosophila species, *D. yakuba* and *D. erecta,* we observe seven amino acid substitutions along the lineage leading to *D. melanogaster* and *D. simulans*. Given this observation, we chose to apply the MKT to these species in a manner similar to its first implementation (McDonald and Kreitman 1991) by combining the polymorphism from *D. melanogaster* and *D. simulans*. Our divergent changes in this MKT comparison are all differences since the most recent common ancestor of the *D. melanogaster* and *D. simulans* lineage rooted by the *D. yakuba* and *D. erecta* lineages. This combined polymorphism MKT rejects neutral expectations, in the direction suggestive of an excess of amino acid substitutions.

**Table 2.** Results of the McDonald-Kreitman Test at *Sxl* between 3 different sets of *Drosophila* species.

|  | Synonymous | Nonsynonymous | *P*-value <sup>a</sup> |
| --- | --- | --- | --- |
| Single species polymorphism |  |  |  |
| <u>*D. ananassae/D. atripex*</u> |  |  |  |
| polymorphic | 8 | 0 |  |
| Fixed divergent | 44 | 2 | 1.00 |
| <u>*D. pseudoobscura/D. miranda*</u> |  |  |  |
| polymorphic | 9 | 1 |  |
| Fixed divergent | 14 | 0 | 0.417 |
| Two species polymorphism combined |  |  |  |
| <u>*(D. melanogaster+D. simulans)/D. yakuba*</u> |  |  |  |
| polymorphic | 19 | 0 |  |
| Fixed divergent | 19 | 7 | **0.015** |

<sup>a</sup> Fisher exact test *P*-value

The significant MKT for *D. melanogaster/D. simulans* could be due to selective fixation of amino acid differences or to selection acting on synonymous changes. While synonymous sites have traditionally been assumed as the neutral yardstick of molecular evolution, there is evidence that this assumption may be invalid, for at least some genes in *Drosophila* (e.g., (Bauer DuMont et al. 2004; Bauer DuMont et al. 2009; Poh et al. 2012; Lawrie et al. 2013). We looked for evidence of selection acting on synonymous changes at *Sxl* using a per site counting method (CF-test) similar to a dN/dS comparison (Bauer DuMont et al. 2004), except in this test we are comparing the number of changes toward unpreferred or preferred codons per the number of unpreferred and preferred “sites”. Along the *D. simulans*, *D. ananassae* and *D. pseudoobscura* lineages we observe a significant departure from neutrality in the direction suggesting a selective advantage of mutations toward preferred codons at *Sxl* (Table 3). The test did not reject in *D. melanogaster*.

**Table 3.** Results of the CF Test at *Sxl* along four *Drosophila* lineages.

|  | Unpreferred | Preferred | *P*-value <sup>a</sup> | direction of departure |
| --- | --- | --- | --- | --- |
| ***D. melanogaster* lineage** |  |  |  |  |
| Fixed substitutions | 5 | 0 |  |  |
| Sites | 150 | 31 | 0.303 |  |
| ***D. simulans* lineage** |  |  |  |  |
| Fixed substitutions | 1 | 2 |  |  |
| Sites | 150 | 31 | 0.022 | preferred codons favored |
| ***D. ananassae* lineage** |  |  |  |  |
| Fixed substitutions | 6 | 11 |  |  |
| Sites | 174 | 29 | <0.001 | preferred codons favored |
| ***D. pseudoobscura* lineage** |  |  |  |  |
| Fixed substitutions | 1 | 3 |  |  |
| Sites | 154 | 31 | 0.002 | preferred codons favored |

<sup>a</sup> Fisher exact test *P*-value

Selection acting on synonymous sites can lead to false positive MKT results by elevating the synonymous polymorphism cell of the 2×2 table, due to segregating slightly deleterious synonymous variants. One method proposed to mitigate the effects of selective constraint on the MKT is to remove low frequency polymorphisms from the analysis (Fay et al. 2002). Given the CF-test results for *D. simulans*, we also carried out the MKT removing two derived unpreferred polymorphic sites at low frequency (singletons – with a frequency of 10% in sample) in *D. simulans*. The *D. melanogaster/D. simulans* MKT remains significant (*P*-value = 0.031).

To further assess the molecular evolution at *Sxl* we made a coding sequence (CDS) alignment using the program Muscle for *Sxl’s* Isoform L (6 exons - the female specific splice variant; Bell et al 1988) for the following 20 Drosophila species: *D. melanogaster, D. sechellia, D. yakuba, D. erecta, D. eugracilis, D. ficusphila, D. rhopaloa, D. elegans, D. takahashii, D. biarmipes, D. kikkawae, D. bipectinata, D. ananassae, D. miranda, D. pseudoobsura, D. persimilis, D. willistoni, D. mojavensis, D. virilis, and D. grimshawi*. We observe 86 amino acid substitutions at 53 codon positions among these species at *Sxl*. Roughly 38% of the codons that have experienced an amino acid substitution have been hit multiple times (20/53; note that some multiply hit amino acid positions had more than three different amino acids segregating among the species). The conservation of the RNA binding domain of the Sxl protein has been previously noted (Zhang, Klein, Nei 2013). In agreement, 84 out of the 86 amino acid changes occurred outside the RNA binding domain region, between codons 1 - 136 and 304 – 373 in our alignment (Figures 1 and 2). We will call these non-RNA binding regions of the Sxl protein the N-terminal and C-terminal regions, respectively. The amino acid substitutions at Sxl have not occurred equally between the N-terminal and C-terminal regions after taking into account their differences in total codon length. We observe significantly more amino acid substitutions in the C-terminal region (24 codons with an amino acid substitution out of 137 total codons in N-terminal region versus 27 substituted codons out of 70 total codons in C-terminal region; 2×2 table chi-square = 11.1, *P*-value = 0.001). The two regions have experienced a similar proportion of multiple hit codons (10 multiple hit codons out of 24 total codons with an amino acid substitution in N-terminal region versus 10 out of 27 such codons in C-terminal region).

**Figure 1.**
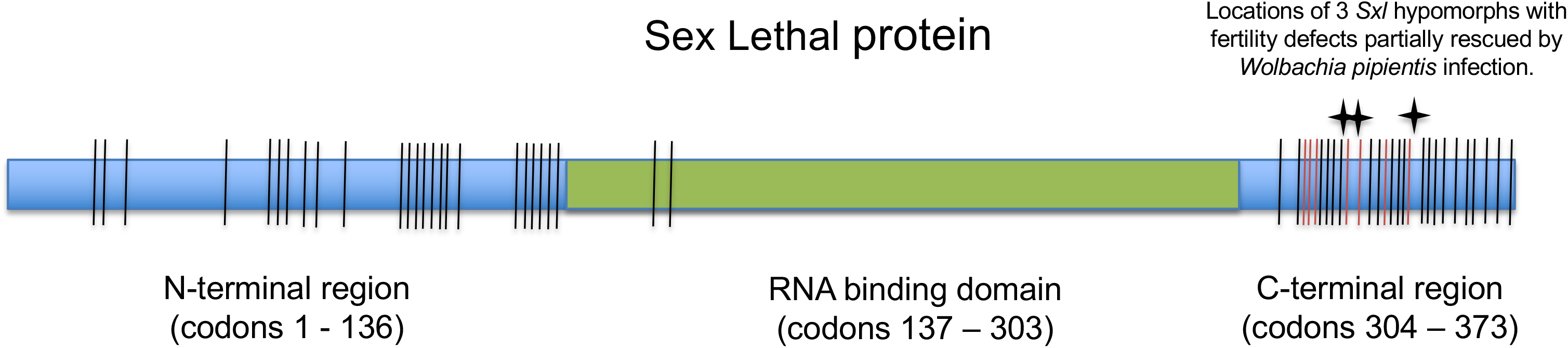
Schematic of Sxl protein. Green box denotes the location of the RNA binding domain of the Sxl protein. Vertical lines show the locations of all amino acid substitutions at *Sxl* across 20 Drosophila species with the red lines being those that have occurred specifically on the lineage leading to *D. melanogaster* and *D. sechellia*. Stars at C-terminal end of protein denote the relative location of the mutations that generate the mutant alleles shown to interact genetically with *Wolbachia pipientis*.

**Figure 2.**
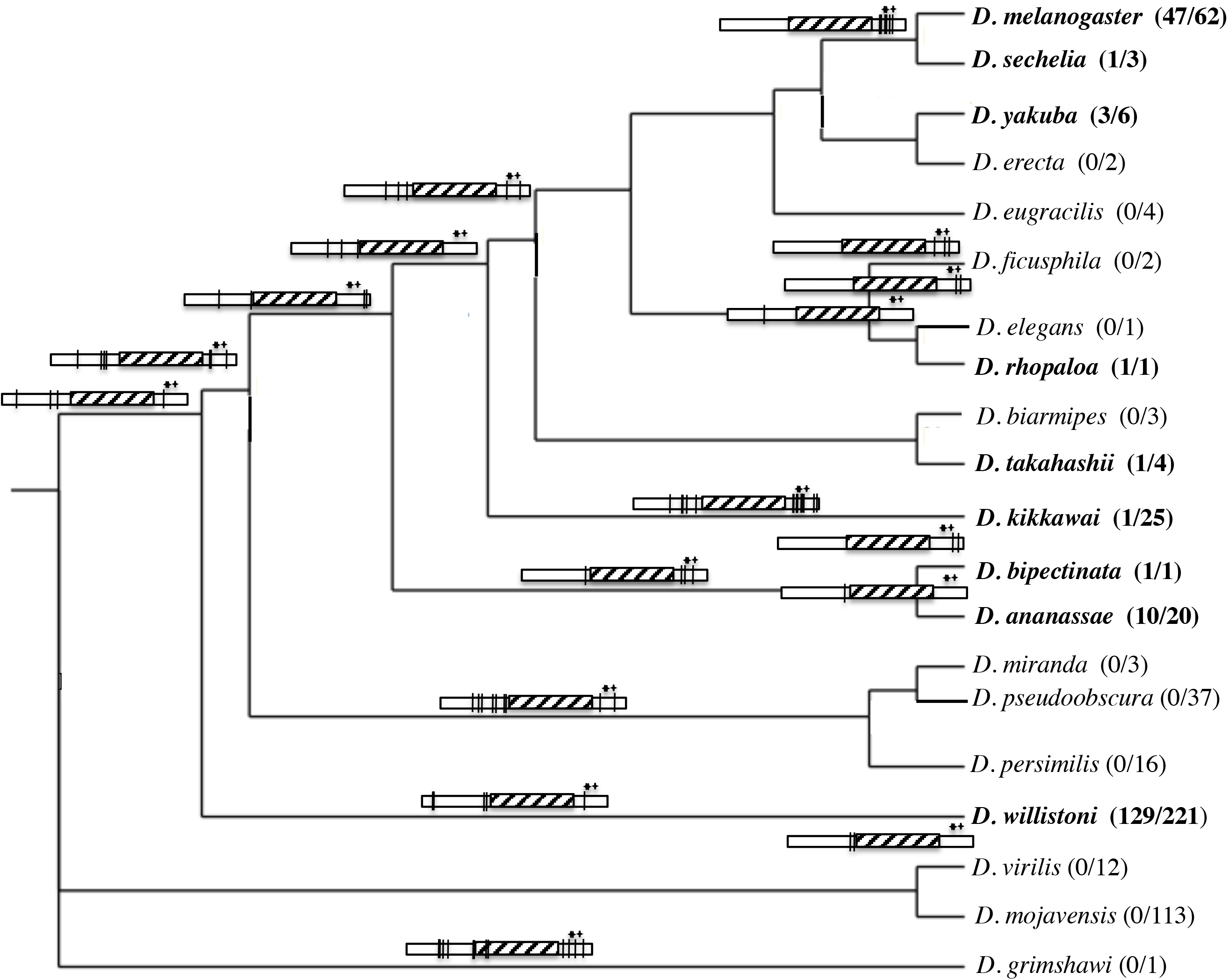
*Sxl* gene tree schematically showing the branches for which amino acid substitutions have occurred. The rectangles denote the Sxl protein with the vertical black lines indicating the location of the amino acid change(s) along that lineage. Hatched box denotes the location of the RNA binding domains of the Sxl protein. Stars at C-terminal end of protein denote the relative location of the three mutations that generate the mutant alleles shown to interact genetically with *W. pipientis*. Species names in bold indicate *Wolbachia* has been detected with the numbers of *Wolbachia* positive lines relative the total number of lines screened given in parenthesis. *Wolbachia* data for all species are from Mateo et al. (2006), as well as for *D. erecta* (Zabalou 2004), *D. kikkawai* (Bennett et al. 2012), *D. bipectinata* (Ravikumar et al. 2011), and *D. willistoni* (Muller et al. 2013).

To determine if the apparent heterogeneity in amino acid substitution at *Sxl* is the result of positive selection, we analyzed the data using the phylogeny-based Hyphy method (Kosakovsky Pond et al. 2005; Delport et al. 2010). Phylogenetic incongruence between species along a gene sequence, due to sorting of ancestral polymorphisms, can have adverse effects on phylogenetic based inferences of positive selection (Wong et al. 2007). We first performed the Hyphy GARD method (Kosakovsky Pond et al. 2006b), which is designed to detect such incongruences. None were detected at a *P*-value cutoff of 0.10. Therefore, we used a maximum-likelihood gene tree, made from 3^rd^ codon positions, in subsequent analyzes.

We applied the following methods of the Hyphy package to our data: GA-branch (Kosakovsky Pond and Frost 2005), BranchRel (Kosakovsky Pond et al. 2011), MEME (Murrell et al. 2012) and FUBAR (Murrell et al. 2013). GAbranch detects significant heterogeneity across the *Sxl* phylogeny in the rate of nonsynonymous compared to synonymous evolution (the dN/dS ratio). The best fitting model includes 3 rate classes, yet the dN/dS for the highest class is only 0.115 across the *Sxl* locus. The posterior probabilities suggest that the following branches are within the highest dN/dS rate class and that they are evolving at a significantly different rate than other branches in the tree: the branch leading to *D. melanogaster* and *D. sechellia*, the branch leading to *D. elegans* and *D. rhopaloa*, the branch leading to the melanogaster species group, and the *D. kikkawei* lineage (Figure 2).

The BranchRel method pools information across sites to estimate selection parameters along branches. The method reports the proportion of codons along each lineage that have evolved under three selection regimes: negative selection, neutral/nearly neutral or episodic positive selection. BranchRel confirmed ubiquitous evidence of amino acid constraint (negative selection) across the *Sxl* phylogeny. After multiple testing corrections, no lineage has significant evidence of positive diversifying selection. Similar results were obtained using MEME.

FUBAR is used to detect selection pressure acting on individual codons. The strength of this method is that it does not restrict the parameter space for which nonsynonymous and synonymous rates are drawn from during the maximum likelihood process. FUBAR does not detect any sites under diversifying selection with a posterior probability greater than 0.90. However, it does detect 258 codons under negative selection with a posterior probability greater than 0.90, which is roughly 70% of the protein. Just over half of these negatively selected sites (140) are located within the RNA binding domain. We observe no significant difference in the number of negatively selected sites between the N-terminal and C-terminal regions of *Sxl* relative to their respective lengths (N-terminal: 77 negative selective sites in 137 codons versus C-terminal: 41 negative selected sites in 70 codons; 2×2 chi-square = 0.023, *P*-value = 0.879).

Interestingly, the seven amino acid differences observed on the lineage leading to the ancestor of the *D. melanogaster* and *D. simulans* species group (with which we observe the significant MKT), cluster with the location of the mutations previously shown to genetically interact with *W. pipientis* (Figure 2).

## Discussion

In this study we use population and phylogenetic based methods to examine the molecular population genetics and evolution of the *Sxl* locus within the genus *Drosophila* for which *Sxl* is the master switch in sex-determination. We were motivated by the observation of a genetic interaction between *W. pipientis* infection and some mutant alleles at *Sxl* in *D. melanogaster* (Starr and Cline 2002; Sun and Cline 2009). It is not known if this interaction is due to a ubiquitous effect of *W. pipientis* on overall egg production, or if it is due to a direct interaction between the endosymbiont and the *Sxl* locus or protein product.

Considering polymorphism data alone, we detect patterns consistent with a recent selective sweep in both *D. simulans* and *D. melanogaster* with the Tajima *D* and Fay and Wu *H* tests, respectively. These significant skews in the frequency spectrum in these species could be due to a variety of evolutionary forces including demography, a selective sweep associated with the fixation of a linked positively selected mutation, or segregating weakly deleterious mutations. *D. simulans* is thought to have experienced a recent population expansion resulting in a general tendency for loci in this species to have negative Tajima *D* test statistics (Kliman et al. 2000). The significant Fay and Wu’s *H* test in *D. melanogaster* remains significant even when demography is incorporated into the null distribution of the test, suggesting that in this species we are detecting a recent selective sweep. These signatures of positive selection would not be due to an amino acid fixation, given that there are no amino acid differences between *D. melanogaster* and *D. simulans* at *Sxl*.

*Sxl’s* long-term evolution within *Drosophila* largely reflects strong conservation of protein sequence, as noted previously by Mullon et al. (2012). We detect the action of negative selection both along lineages and at specific codons. The FUBAR method estimates that 70% of the *Sxl* codons are selectively constrained, suggesting that negative selection has had a pervasive effect on *Sxl’s* molecular evolution. However, we do observe amino acid differences across these *Drosophila* species and the pattern of these substitutions is heterogeneous. For example, GA-branch method detects significant variation in the dN/dS ratio across the *Drosophila* species included in this analysis. This heterogeneity at *Sxl* could be due to sporadic relaxations of the negative selection that dominates *Sxl’s* molecular evolution or due to sporadic bursts of positive selection.

The Hyphy methods used to detect recurring or episodic positive selection fail to do so after multiple testing corrections. However, we note that the pervasive negative selection observed at *Sxl* could confound these methods, especially if the positive selection is weak or if only a few sites are affected (Kosakovsky Pond et al. 2011). Our current data shows no evidence of long-term or recent positive selection at *Sxl* along the *D. ananassae* and *D pseudoobscura* lineages. In contrast, there are weak signatures of both types of selection in *D. melanogaster* and *D. simulans*, so we focus on the molecular evolution of these species and the lineage leading to their common ancestor.

The lineage leading to *D. melanogaster and D. simulans* shows a decoupling of synonymous and nonsynonymous evolution with the MKT in the direction of an excess of nonsynonymous divergence. This result is due to seven amino acid substitutions on the lineage leading to the *D. melanogaster/D. simulans* clade. These nonsynonymous substitutions cluster with the locations of the *Sxl* alleles that genetically interact with *W. pipientis.* In addition, this region of the Sxl protein (the C-terminal non-RNA binding region) has experienced significantly more amino acid substitutions than the N-terminal region. This elevation in amino acid substitutions does not appear to be due to a simple relaxation of constraint as we observe no difference between the C- and N-terminal regions in the number of codons predicted to be experiencing negative selection. These results are suggestive of positive selection being at least partially responsible for the fixations of these seven *D. melanogaster/D. simulans* amino acid substitutions. However, there are other possible explanations for these results such as synonymous site evolution and/or changes in effective population. The seven amino acid fixations could be due to the fixation of slightly deleterious mutations if the effective population size was smaller on the branch leading to the *D. melanogaster/D. simulans*. However, the relaxation of constraint is expected to affect both synonymous and nonsynonymous substitutions. We assume the ancestral state in codon preference is toward preferred synonymous codons at *Sxl*, given the results of the CF-test. If relaxation of constraint were responsible for the burst of amino acid fixations on the *D. melanogaster/D. simulans* lineage we may also expect a burst of derived unpreferred substitutions, but we do not observe this. We observe an equal number of preferred and unpreferrred changes on this lineage (data not shown).

Infection dynamics of *W. pipientis* appear to be sporadic and variable both between and within lineages, with uninfected species interspersed with infected species throughout the phylogeny (e.g., Mateos et al. 2006) consistent with multiple losses or gains of infection. The resulting uncertainty in the infection history of species unfortunately prevents us from reliably testing for correlations between *W. pipientis* infection status and burst of positive selection; there are many factors that could weaken our ability to detect an association.

In this study we present data revealing both similarities and difference between the molecular evolution at *bam* and *Sxl*, two loci that genetically interact with *W. pipientis* infection. For both loci the fixation of amino acid variants appears to be heterogeneous, potentially weakening phylogenetic methods to detect positive selection or associations with character states. The MKT does reject neutrality for both loci in a manner suggestive of an acceleration of nonsynonymous fixations. However, the extent of amino acid differences is very different between these loci with there being 59 fixed amino acid substitutions between *D. melanogaster* and *D. simulans* at *bam* and none at *Sxl*.

Our results do not allow us to draw strong conclusions regarding the role of positive selection on the molecular evolution of the *Sxl* locus within *Drosophila*, but it also does not allow us to discount the influence of both long term (MKT) or recent (Tajima D and Fay and Wu H tests) positive selection in *D. melanogaster* and *D. simulans*. Current data and methodology also do not allow us to make a direct connection between *W. pipientis* infection and selective pressures acting on *Sxl* or *bam*. So, it remains open whether the genetic interaction between mutant *Sxl* and *bam* alleles and *W. pipientis* is due to a direct interaction between Sxl and Bam protein and this endosymbiont. Our results do motivate screening for genetic interactions between *W. pipientis* and other mutant alleles at *Sxl* and other GSC loci because the observation that *W. pipientis* rescues some but not other mutations (e.g., Starr and Cline 2002) will help refine candidates for the mechanism(s) by which *W. pipientis* is manipulating *Drosophila* reproduction.

## Acknowledgements

We would like to thank Jae Young Choi for helpful input regarding data collection and manuscript preparation, and Helen K. Salz for her feedback on the manuscript. Research was supported by National Institute of Health grant number R01GM095793 to C.F.A.

